# Striking reduction in neurons and glial cells in anterior thalamic nuclei of older patients with Down’s syndrome

**DOI:** 10.1101/449678

**Authors:** James C. Perry, Bente Pakkenberg, Seralynne D. Vann

## Abstract

The anterior thalamic nuclei are important for spatial and episodic memory; however, there is surprisingly little information about how these nuclei are affected in many conditions that present with memory impairments, including Down’s syndrome. To assess the status of the anterior thalamic nuclei in Down’s syndrome we quantified neurons and glial cells in the brains from four older patients with this condition. There was a striking reduction in the volume of the anterior thalamic nuclei and this appeared to reflect the loss of approximately 70% of neurons. The number of glial cells was also reduced but to a lesser degree than neurons. The anterior thalamic nuclei appear to be particularly sensitive to effects of aging in Down’s syndrome and the pathology in this region likely contributes to the memory impairments observed. These findings re-affirm the importance of assessing the status of the anterior thalamic nuclei in conditions where memory impairments have been principally assigned to pathology in the medial temporal lobe.

## 1. Introduction

Down’s syndrome (DS; trisomy 21) is the most common chromosomal disorder, affecting 1 in 1000 live births in the UK (Wu and Morris, 2013). A common feature of DS is learning disabilities with accompanying language and memory impairments. Over recent years there has been a noticeable increase in life expectancy for adults with DS, with the median life expectancy in England and Wales currently at 58 years (Wu and Morris, 2013). This increase in lifespan has resulted in a concomitant increase in the number of adults with DS affected by dementia, particularly Alzheimer’s disease, which is considerably more prevalent in adults with DS than the general population (e.g. Zigman and Lott, 2007). Given the increased longevity of DS patients, it has become increasingly important to understand the underlying changes that occur with age and dementia in this condition.

Research into mnemonic impairments in DS has typically focused on medial temporal lobe structures, which are smaller in DS patients even when overall brain size is taken into account (Carducci et al., 2013; Kemper 1991; Krasuski et al., 2002; Pinter et al., 2001). However, there is an increasing realisation that it is necessary to look beyond the hippocampal formation in conditions that present with memory impairments. In doing so, the anterior thalamic nuclei (ATN) become an obvious target due to their connections with the hippocampal formation and their importance for memory (Dillingham et al., 2015; Harding et al., 2000; Jankowski et al., 2013; Van der Werf et al., 2003). The presence of neurofibrillary tangles and plaques in the ATN of patients with Alzheimer’s disease further highlights the potential relevance of this structure to DS (Aggleton et al., 2016; Braak and Braak, 1991; Rub et al., 2016). There is very little currently known about the ATN in patients with DS as neuroimaging studies have only focused on the thalamus as a whole (e.g. Annus et al., 2017; Teipel et al., 2004). There are limitations in treating the thalamus as a unitary structure given the many differences in connectivity and function of the various nuclei. Due to their size and location, it is difficult to obtain accurate measures of ATN volume using current neuroimaging techniques, which makes post-mortem assessments all the more important. Moreover, post-mortem tissue enables assessments at the cellular level in addition to volumetric measures. Therefore, stereological counts of neurons and glial cells were made in the ATN of four aged female brains with DS and six age-matched controls.

## 2. Method

### 2.1 Patient characteristics

Brains from four female Down’s syndrome (DS) patients (mean age 69, range, 61 – 80) were provided by the Netherlands Brain Bank and six female age-matched controls (mean age 71, range, 60 – 85) were obtained post-mortem according to Danish law regarding the use of autopsied human tissue in research. The diagnosis of DS was based on the characteristic physical appearance of patients and presence of intellectual disability as post-mortem chromosomal analysis was not available (see Table 1 for clinical data). Background details and neuropathological information will be provided for each patient in turn:

Patient 1 was first noted as having seizures at the age of 36 with an EEG showing mild diffuse cerebral disorder with focal abnormalities in both temporal regions. At this point the patient was treated with anti-epileptic medication. At the age of 53 the patient had been free from seizures for many years but started showing the first symptoms of dementia. Two years following the onset of dementia symptoms, Patient 1 was admitted to a nursing home where she gradually deteriorated both mentally and physically. A year prior to death she suffered from several epileptic seizures. The post-mortem found there to be numerous senile plaques and neurofibrillary tangles in a disrupted interneuronal network within the frontal, temporal, parietal and occipital lobes and the hippocampus. The substantia nigra showed very little if any cell loss and the pons, medulla oblongata and cerebellum all appeared normal.

Patient 2 functioned at a moderate level for the majority of her life, working in a social working facility for almost 30 years until the age of 50. At the age of 57 she was admitted to a home due to increasing problems in taking care of herself, often getting lost, suffering from mood swings and having difficulty taking care of personal needs. At the age of 60 it was thought she had between stage 3 and stage 4 Alzheimer’s disease and she started to have minor epileptic insults. EEG showed a diffuse abnormality. According to the post-mortem neuropathology report the brain was severely atrophied with numerous plaques and tangles in the interneuronal network across all lobes and in the hippocampus. The pons, putamen, medulla oblongata, and cerebellum all showed no abnormality.

Patient 3 was diagnosed with epilepsy at the age of 26. An EEG carried out when the patient was 37 showed diffuse irregular activity. She was admitted into a home at the age of 44. At the age of 60 the patient had a marked decrease in general functioning with the likely diagnosis Alzheimer’s disease and at 64 she was thought to have stage 2 Alzheimer’s disease. A report from the following year stated that she was unable to dissociate between members of her family and nursing home staff, she was disoriented in time and place with difficulties in communication. In the last two months before death a cognitive assessment showed her linguistic and communicative skills to be severely disturbed as were her memory function, orientation and concentration. The post-mortem report identified numerous tangles and plaques in the cortex, hippocampus and amygdala. The cingulate gyrus showed severe spongiosis while the medulla oblongata, medulla cervicalis, and cerebellum showed no abnormalities.

Patient 4 was admitted to a nursing home at the age of 66 due to a diagnosis of dementia. However, the patient remained alert until her death and continued to recognise people and make demands. An EEG carried out when the patient was 70 showed decreased alpha rhythm and an epileptic predisposition although clinically there was no indication of epilepsy. Post-mortem revealed numerous plaques and tangles in a disrupted interneuronal network across the cortex and hippocampus. The substantia nigra, cingulate gyrus, locus coeruleus, pons and cerebellum showed no abnormalities. `The amygdala showed only a few neurofibrillary tangles and senile plaques.

### 2.2 Tissue Processing

Brains were fixed with 0.1M sodium phosphate buffered 4% formaldehyde (pH 7.2) for at least five months. The cerebrum was detached at the level of the third cranial nerve, the meninges were removed and hemispheres were processed as previously described (Karlsen and Pakkenberg, 2011). Coronal sections 40µm thick were stained with Giemsa’s azur eosin methylene blue solution (Merck, Darmstadt, Germany) and KHPO4 (pH 4.5) in a 1:4 ratio. Slides were coded by a researcher not involved in data collection to allow blinded analyses.

**Table 1.** Patient clinical details

*Patient clinical details*
|  | Age (years) | Height (cm) | Hemisphere (g) | Country | Cause of death | PMD (h) |
| --- | --- | --- | --- | --- | --- | --- |
| Controls | 75 | 159 | 575 | DK | AMI | 24:00 |
|  | 85 | 159 | 590 | DK | AMI | 24:00 |
|  | 64 | 154 | 620 | DK | AMI | 24:00 |
|  | 60 | 161 | 520 | DK | AMI | 48:00 |
|  | 70 | 169 | 667 | DK | AMI | 24:00 |
|  | 74 | 160 | 590 | DK | AMI | 24:00 |
| Mean | 71.3 | 160.3 | 594 |  |  |  |
| DS | 61 | - | 444 | NED | Pneumonia | 05:30 |
|  | 64 | 148 | 371 | NED | Pneumonia | 04:00 |
|  | 70 | 162 | 387 | NED | Ileus | 09:00 |
|  | 80 | - | 537 | NED | AMI/pneumonia | 04:00 |
| Mean | 68.8 | 155 | 435 |  |  |  |

Note: AMI, acute myocardial infarction; DK, Denmark; DS, Down's Syndrome; NED, The Netherlands; PMD, post-mortem delay

### 2.3 Cell number and volume estimation

Due to the difficulty in distinguishing between the anteromedial and anteroventral thalamic nuclei in Nissl-stained human tissue, these nuclei were grouped together as the anterior principal nuclei (APn; Fig. 1) and delineated as previously described (Armstrong, 1990; Harding et al., 2000; Hirai et al., 1989). The APn is in the main encapsulated by fibres making it easily identifiable; only the inferior edge of the structure does not have a clear boundary. The ventral medial portion of the APn was distinguishable from the anterodorsal, paratenial, and paraventricular thalamic nuclei, and the stria medullaris of thalamus by lamina encapsulating these nuclei, making the APn discernible from these midline nuclei. There was a maximum of one section per brain where the central medial thalamic nuclei were not as distinct from the ventral medial tip of the APn. For these sections, where possible, the lamina separating the paraventricular and central medial thalamic nuclei from the APn was used as a guide for distinguishing the ventral medial portion.

The number of neurons and glial cells were estimated from one hemisphere per brain using the optical fractionator method (Gundersen, 1986; West et al., 1991). Slides were mounted on an Olympus BX60 microscope equipped with a computer driven X – Y motorised stage (ProScan III Prior, UK) with Z-axis focus control (Heidenhain Encoder, Germany) and a CMOS camera (Basler, Germany) connected to a computer running the NewCast Stereology software package (Visiopharm, Hørsholm, Denmark). Differentiation of neuron and glial cell types was based on cell morphology as previously described (Karlsen and Pakkenberg, 2011; Pelvig et al., 2008). In brief, neurons have a single large nucleolus and a large nuclei surrounded by a visible cytoplasm. Neurons can be further classified into ‘small’ local inhibitory neurons and ‘large’ projecting excitatory neurons based on size (Armstrong, 1990; Dorph-Petersen et al., 2004). These two types of neurons are only referred to as small and large in this study as definitive classification of neurons cannot be achieved with Nissl staining alone (Hou et al., 2012; Lyck et al., 2006). Oligodendrocytes were identified as having a small rounded nucleus with dense chromatin and a perinuclear halo. Astrocytes have a pale nucleus surrounded by a translucent cytoplasm and heterochromatin concentrated in granules in a rim below the nuclear membrane. Microglia appear small and elongated or comma shaped with dense peripheral chromatin. The nucleus was the counting item for both neurons and glial cells during sampling.

**Figure 1.**
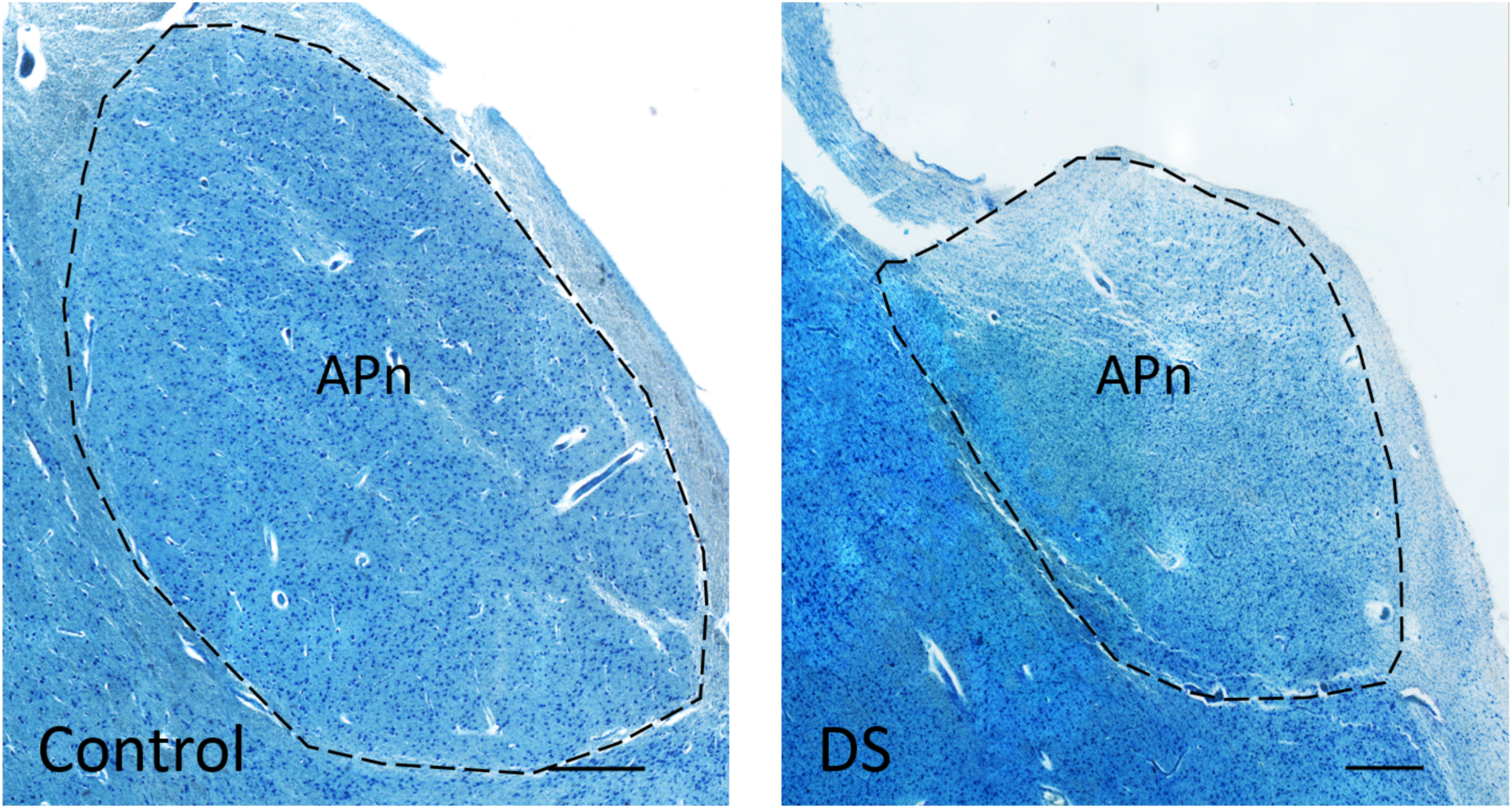
The anterior principal thalamic nucleus (APn) in a control and a Down’s syndrome (DS) brain. Sections were stained with Giemsa’s azur eosin methylene blue solution. The APn is encapsulated by the internal medullary lamina. Image montages of the APn were captured at similar levels using the same magnification. Scale bars = 750µm.

Sampling of sections followed a systematic, uniform random sampling scheme to ensure all parts of the APn had an equal chance of being sampled. A section sampling fraction of 1/16 was employed resulting in 9 – 15 sections sampled per brain (mounted section thickness 40 µm). The contour of the APn was traced live using a 10x objective (0.4 NA) and cells were counted using a 100x oil immersion objective (NA 1.35) with a final onscreen magnification of 3000x. A disector height of 20µm with a 5µm guard zone was used to sample in the z-axis. The counting frame area was 2800 µm^2^ for neurons and 1350 µm^2^ for glial cells. The x – y step length was adjusted to allow optimal sampling of brains considered ‘large’ and ‘small’ but did not change within cases. The x – y step length was set to 400 – 1000 µm^2^ for neurons and 800 – 1000 µm^2^ for glial cells. These sampling parameters led to a mean of 206 neurons (range, 152 – 276) and 631 glial cells
(range, 399 – 1081) counted per brain in a mean of 222 counting frames (range, 131 – 466) for neurons and 103 counting frames (range, 45 – 142) for glial cells. The numbers of neurons and glial cells were estimated from

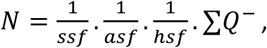

where *ssf* is the section sampling fraction; the area sampling fraction (*asf*), is the ratio of the counting frame area to the area of the x – y step length; the height sampling fraction (*hsf*), is the ratio of the disector probe height to the Q^-^ weighted tissue thickness; and ∑ Q^-^ is the total count of cells sampled (Dorph-Petersen et al., 2004). The density of neurons and glia was calculated as N/volume.

The unilateral volume of the APn was estimated according to Cavaleri’s principle (Gundersen and Jensen, 1987; Gundersen et al., 1999) but was not corrected for tissue shrinkage as the degree of shrinkage between control and DS brains was estimated to be the same in a previous study using the same samples (Karlsen and Pakkenberg, 2011).

The coefficient of error (CE) was calculated (Gundersen et al., 1999) to provide information regarding the precision of the estimates. The CE for group estimates of each measure was estimated as:

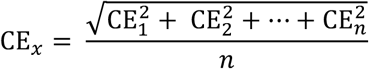

Where CE*_x_* is the mean CE estimate and *n* is the number of individuals. As a measure of the variability of the estimates the observed inter-individual coefficient of variation (CV = standard deviation/mean) are reported in parentheses after group means in Fig. 1.

### 2.4 Statistics

Differences between groups were tested using an independent means *t*-test (two-tailed). Welch’s *t*-test for unequal variances was used where appropriate. A Mann-Whitney U test was used when normality tests failed. Where relevant, effect sizes between groups were expressed as Hedges *g* when equal variances were assumed, Glass’s Δ when equal variances were not assumed, or η*^2^*following a Mann-Whitney U test (Fritz et al., 2012). The common language effect size (*CL*), the percentage of occasions that a randomly sampled case from distribution with the higher mean will have a higher score compared to a randomly sampled case from the distribution with the lower mean (Lakens, 2013; McGraw and Wong, 1992), is also reported for the key measures total neurons, total glial cells, and volume to conceptualise the size of these effects. SPSS software (version 20, IBM Corporation) was used to carry out statistical analyses. The Holm-Bonferroni procedure was used to correct for multiple comparisons. The corrected *P-*values (*P*_c_) of less than 0.05 were considered significant.

The CE estimates for the total number of neurons was 0.07 for both control and DS brains, for large neurons was 0.09 for both groups, and for small neurons was 0.14 for control and 0.11 for DS brains. The CE estimates for total glial cells was 0.04 for both control and DS brains, for oligodendrocytes was 0.07 for control and 0.08 for DS brains, for astroglia was 0.05 for both groups, and for microglia was 0.51 for controls and 0.49 for DS brains. Estimates of final CE for volume was 0.05 both control and DS brains.

## 3. Results

All estimates of anterior principal thalamic nuclei (APn) neurons, glial cells, volume, and density are reported as unilateral numbers, i.e. measures for a single hemisphere, and are shown in Fig. 2.

As shown in Fig. 2, there was a 68% reduction in total numbers of neurons in the APn of the Down’s syndrome (DS) brains compared to controls. This difference was found to be significant (*t*(7.98) = 6.99, *P*_c_ < .001; Δ = 3.68, 95% CI 1.67 – 5.81, *CL* = 99%; Fig. 2A). This overall reduction reflected changes in both large (*t*(8) = 5.26, *P*_c_ < 0.01; *g* = 3.06, 95% CI 1.22 – 4.91; Fig. 2B) and small neurons (Mann-Whitney U = 0.00, *P*_c_ = 0.029; η^2^ = 0.65; Fig. 2C). There was a significant 37% reduction in estimated total number of glial cells in DS brains compared to controls (*t*(8) = 3.43, *P*_c_ = 0.036; *g* = 2.00, 95% CI 1.48 – 5.43, *CL* = 93%; Fig. 2D). A significant reduction in DS brains was also observed when glial cells were classified as astroglia (t(8) = 3.58, *P*_c_ = 0.036; g = 2.09, 95% CI 0.53 – 3.65; Fig. 2E). The difference for oligodendrocytes was borderline significant when corrected (*t*(8) = 2.73 *P*_c_ = 0.052, *P* = 0.026; g = 1.59, 95% CI 0.14 – 3.03; Fig. 2F). The estimated volume of the APn was 62% less in DS brains than controls and this reduction was significant (*t*(7.99) = 10.09, *P*_c_ < .001; Δ = 5.27, 95% CI 2.70 – 8.05, *CL* = 99%; Fig. 2G). The estimated density of total neurons for DS brains did not differ from the estimate for controls (t(8) = 1.33, *P* = 0.22; Fig. 2H). However, the estimated total glial cell density in DS brains was significantly higher than the estimate in controls, (*t*(8) = 4.12, *P*_c_ = 0.02; *g* = 2.40, 95% CI 0.76 – 4.05; Fig. 2I). For microglia, the number available for sampling was consistently low (2 – 20 per brain), resulting in high group CE values (0.49 – 0.51). Controls were estimated to have 1.64 x 10^6^ (CV, 0.93) microglia whereas DS brains had 0.94 x 10^6^ (CV, 0.51) microglia, although this reduction was not significant (*P* > 0.40).

**Figure 2.**
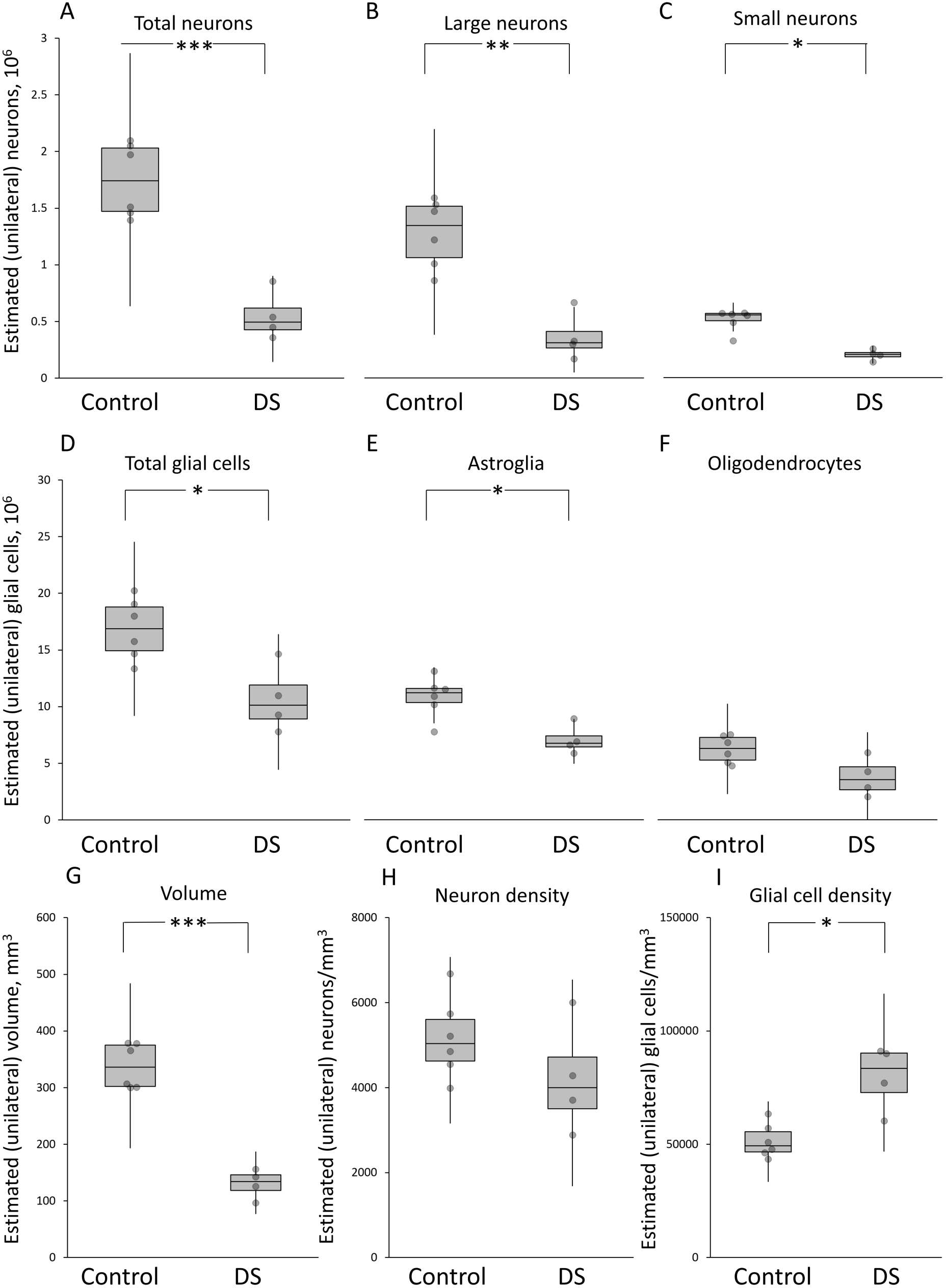
Unilateral estimates for anterior principle thalamic nucleus (APn) in Control and Down’s Syndrome (DS) brains. Top panel, (**A**) total neurons (combined large and small neurons; control = 1.75 x 10^6^ (0.19), DS = 0.55 x 10^6^ (0.39)), (**B**) large (‘projecting’) neurons (control = 1.25 x 10^6^ (0.23), DS = 0.36 x 10^6^ (0.58)), and (**C**) small (‘local inhibitory’) neurons (control = 0.50 x 10^6^ (0.19), DS = 0.20 x 10^6^ (0.24)). Middle panel, (**D**) total glial cells (combined astroglia, oligodendrocytes, and microglia; control = 16.84 x 10^6^ (0.16), DS = 10.68 x 10^6^ (0.28)), (**E**) astroglia (control = 10.60 x 10^6^ (0.17), DS = 6.92 x 10^6^ (0.18)), and (**F**) oligodendrocytes (Control = 6.08 x 10^6^ (0.19), DS = 3.67 x 10^6^ (0.45)). Bottom panel, (**G**) volume of APn (control = 338 mm^3^ (0.12), DS = 130 mm^3^ (0.20)), (**H**) neuron density (control = 5171 x mm^3^ (0.18), DS = 4221 x mm^3^ (0.31)), and (**I**) glial cell density (control = 51,509 x mm^3^ (0.14), DS =79,649 x mm^3^ (0.18)). The central line in each box indicates the median value. The box extends from the first to the third quartile range. The whiskers extend 1.5x the interquartile range. Individual data points are shifted along the x-axis to aid visualization of overlapping data points. Note, in (**F**) the whisker for the DS plot extends slightly below zero (not shown). Note, values in this legend are mean estimates with the coefficient of variation reported in parentheses. *Significantly different from Control, *P*_c_ < 0.05; **significantly different from Control, *P*_c_ < 0.01; ***significantly different from Control, *P*_c_ < .001.

## 4. Discussion

We estimated numbers of neurons and glial cells in the combined anteroventral and anteromedial thalamic nuclei (anterior principal thalamic nuclei, APn) of aged women with Down’s syndrome (DS). When compared to age-matched controls, there were striking changes with almost 70% fewer neurons in the DS brains, with reductions in both large and small neurons. Glial cells were also reduced but to a lesser degree (37%), with comparable changes across astrocytes and oligodendrocytes. The overall volume of APn was also markedly reduced (62%) in DS patients. As a result there was no difference in neuronal density between DS patients and controls. A similar pattern is found in Korsakoff patients whereby neuron density is unaffected but overall numbers are reduced by ∼50% (Harding et al., 2000). This suggests that substantial tissue atrophy in the APn is associated with neuronal loss and highlights the importance of assessing overall cell counts and not just neuronal density.

While the current study focused on the APn, previous studies using the same brains have reported changes in the mediodorsal thalamic nucleus and cortical regions (Karlsen et al., 2014; Karlsen and Pakkenberg, 2011). While the volume of the mediodorsal thalamic nucleus was reduced to a similar extent as the APn (59%), the overall cell loss was less with a 43% reduction in neurons but no changes in glial cells (Karlsen et al., 2014). This highlights the limitations of treating the thalamus as a unitary structure. Furthermore, it becomes clear that simple volumetric measures can mask substantial variations in patterns of cell loss. Compared to the cortex, there were greater reductions in the APn when considering both volume and neuronal count reduction (i.e. ∼40% for both in cortex; Karlsen and Pakkenberg, 2011). In contrast, the reduction in neocortical glial cells was more comparable to the current findings (i.e. ∼30%; Karlsen and Pakkenberg, 2011). Together, the changes found in APn are more pronounced than other areas assessed in the same brains, particularly when considering the reduction in neurons.

While there are some differences between the patient and control groups in terms of cause of death and post-mortem delay (see Table 1), the present findings are unlikely to be due to these factors (Blair et al., 2016). The basal ganglia were also examined in these same brains and there was no neuronal loss in this structure (Karlsen and Pakkenberg, 2011), confirming the specificity of the current findings.

An obvious consideration is the extent to which these APn changes simply reflect the accompanying dementia in this patient group rather than being directly related to DS, especially given the finding that neuronal numbers within some thalamic nuclei have been shown to correlate with amyloid (Erskine et al., 2016; Erskine et al., 2017). Pathology related to Alzheimer’s disease (AD) is a common feature in DS patients with the onset of dementia often occurring by the mid-50s (Head et al., 2012). Across the cortex and hippocampal formation, the development patterns of neurofibrillary tangles are comparable across DS and AD groups although there appears to be more widespread and denser deposition of amyloid plaques in DS (Hof et al., 1995). However, Braak found no noticeable difference in AD-related pathology in patients with or without DS in the anterior thalamus (Braak and Braak, 1991). There are surprisingly few studies that have quantitatively assessed the anterior thalamic nuclei (ATN) in Alzheimer’s disease (without accompanying DS) and even fewer that have carried out neuronal counts. While tangles and plaques are present in the APn, it is the anterodorsal thalamic nucleus that is most severely affected (Braak and Braak, 1991; Rub et al., 2016; Xuereb et al., 1991). Consistent with this, Xuereb et al. (1991) found no significant reduction in volume or neuronal counts in the APn of Alzheimer’s disease patients, although there was a significant reduction in neuronal counts in the anterodorsal nucleus. Similarly, Hornberger et al. (2012) found no significant reduction in ATN volume in Alzheimer’s disease patients using either neuroimaging or post-mortem tissue. There also does not seem to be a straightforward link between severity of AD and the status of the APn in the current cohort, (some detail about relationship between Braak stage and measures). It, therefore, appears that the APn pathology in the current cohort does not simply reflect AD-related changes although the earlier onset of AD-like pathology or the greater presence of amyloid plaques in the cortex and hippocampus in DS may exacerbate the cell loss. The current findings show greater similarity to other dementias such as the semantic variant of primary progressive aphasia and fronto-temporal dementia where there is ∼30-40% decrease in ATN volume (Tan et al., 2014).

The current findings raise the question as to whether ATN pathology is a common feature of DS and whether the ATN are also compromised in younger patients with DS. At present, very little is known about the status of the ATN in this patient group. Studies in younger patients have typically only measured the thalamus as a whole and in doing so have not found any difference between patient and control groups (Annus et al., 2017; but see Gunbey et al., 2017; Jernigan et al., 1993; Weis et al., 1991). However, this approach may be masking more selective changes within the ATN. Reports of enlarged 3^rd^ ventricles in younger DS patients could also be indicative of atrophy within the ATN (Raz et al., 1995; Schimmel et al., 2006). Consistent with this, the mammillary bodies were shown to be markedly smaller in adults with DS, moreover, their size significantly correlated with general aptitude (Raz et al., 1995). Given the reductions in mammillary body and hippocampal volume, and the dense connectivity between these structures and the ATN (Bubb et al., 2017), it is possible that the ATN are also compromised at an earlier time-point in DS patients. Pathology within the extended hippocampal-diencephalic network would be consistent with the pattern of memory impairments observed in DS, where there is greater disruption to explicit compared to implicit memory (Vicari et al., 2000). A similar memory profile is found following damage to the mammillary body-thalamic axis (Carlesimo et al., 2007; Tsivilis et al., 2008; Vann et al., 2009).

One limitation of the current study was the small sample sizes, as is often the case with postmortem studies. However, the pattern of cell loss in all DS brains was very consistent, both in the APn as well as previous structures measured (Karlsen et al., 2014; Karlsen and Pakkenberg, 2011). Furthermore, although we had a low sample size, the differences found on key measures was reliable, with large effect sizes. The sample size was, therefore, sufficient to detect the differences reported.

Together, these data show a striking decrease in both the volume and cell number of APn in aged DS patients. The pathology in this region is likely to contribute to memory impairments given the importance of this region for mnemonic processes. These findings re-enforce the need to look beyond the medial temporal lobe when considering neurological conditions that present with memory impairments. There is also a clear need to further assess this structure in future studies of DS to determine the age of onset of ATN pathology in this patient group.

## Acknowledgments

We thank The Netherlands Brain Bank, department of The Netherlands Institute for Neuroscience, for providing the Down’s syndrome brains. We thank Susanne Sørensen for expert technical assistance and Professors Elizabeth Fisher and John Aggleton for helpful discussion.

## Abbreviations

AD: Alzheimer’s disease
APn: anterior principle nuclei
asf: area sampling fraction
ATN: anterior thalamic nuclei
CE: coefficient of error
CV: coefficient of variation
DS: Down’s syndrome
hsf: height sampling fraction
ssf: section sampling fraction

## References

Aggleton, J.P., Pralus, A., Nelson, A.J., Hornberger, M., 2016. Thalamic pathology and memory loss in early Alzheimer’s disease: moving the focus from the medial temporal lobe to Papez circuit. Brain 139(Pt 7), 1877–1890.

Annus, T., Wilson, L.R., Acosta-Cabronero, J., Cardenas-Blanco, A., Hong, Y.T., Fryer, T.D., Coles, J.P., Menon, D.K., Zaman, S.H., Holland, A.J., Nestor, P.J., 2017. The Down syndrome brain in the presence and absence of fibrillar beta-amyloidosis. Neurobiol. Aging 53, 11–19.

Armstrong, E., 1990. The limbic thalamus: anterior and mediodorsal nuclei, in: Paxinos, G. (Ed.) The human nervous system. Academic Press Inc, San Diego, CA, pp. 469–478.

Blair, J.A., Wang, C., Hernandez, D., Siedlak, S.L., Rodgers, M.S., Achar, R.K., Fahmy, L.M., Torres, S.L., Petersen, R.B., Zhu, X., Casadesus, G., Lee, H.G., 2016. Individual Case Analysis of Postmortem Interval Time on Brain Tissue Preservation. PLoS One 11(3), e0151615.

Braak, H., Braak, E., 1991. Alzheimer’s disease affects limbic nuclei of the thalamus. Acta Neuropathol 81(3), 261–268.

Bubb, E.J., Kinnavane, L., Aggleton, J.P., 2017. Hippocampal - diencephalic - cingulate networks for memory and emotion: An anatomical guide. Brain Neurosci Adv 1(1).

Carducci, F., Onorati, P., Condoluci, C., Di Gennaro, G., Quarato, P.P., Pierallini, A., Sara, M., Miano, S., Cornia, R., Albertini, G., 2013. Whole-brain voxel-based morphometry study of children and adolescents with Down syndrome. Funct Neurol 28(1), 19–28.

Carlesimo, G.A., Serra, L., Fadda, L., Cherubini, A., Bozzali, M., Caltagirone, C., 2007. Bilateral damage to the mammillo-thalamic tract impairs recollection but not familiarity in the recognition process: a single case investigation. Neuropsychologia 45(11), 2467–2479.

Dewulf, A., 1971. Anatomy of the normal human thalamus. Elsevier, Amsterdam; London.

Dillingham, C.M., Erichsen, J.T., O’Mara, S.M., Aggleton, J.P., Vann, S.D., 2015. Fornical and nonfornical projections from the rat hippocampal formation to the anterior thalamic nuclei. Hippocampus 25(9), 977–992.

Dorph-Petersen, K.A., Pierri, J.N., Sun, Z., Sampson, A.R., Lewis, D.A., 2004. Stereological analysis of the mediodorsal thalamic nucleus in schizophrenia: volume, neuron number, and cell types. J. Comp. Neurol. 472(4), 449–462.

Erskine, D., Taylor, J.P., Firbank, M.J., Patterson, L., Onofrj, M., O’Brien, J.T., McKeith, I.G., Attems, J., Thomas, A.J., Morris, C.M., Khundakar, A.A., 2016. Changes to the lateral geniculate nucleus in Alzheimer’s disease but not dementia with Lewy bodies. Neuropathol. Appl. Neurobiol. 42(4), 366–376.

Erskine, D., Thomas, A.J., Attems, J., Taylor, J.P., McKeith, I.G., Morris, C.M., Khundakar, A.A., 2017. Specific patterns of neuronal loss in the pulvinar nucleus in dementia with lewy bodies. Mov Disord 32(3), 414–422.

Fritz, C.O., Morris, P.E., Richler, J.J., 2012. Effect size estimates: current use, calculations, and interpretation. J Exp Psychol Gen 141(1), 2–18.

Gunbey, H.P., Bilgici, M.C., Aslan, K., Has, A.C., Ogur, M.G., Alhan, A., Incesu, L., 2017. Structural brain alterations of Down’s syndrome in early childhood evaluation by DTI and volumetric analyses. Eur. Radiol. 27(7), 3013–3021.

Gundersen, H.J., 1986. Stereology of arbitrary particles. A review of unbiased number and size estimators and the presentation of some new ones, in memory of William R. Thompson. J. Microsc. 143(Pt 1), 3–45.

Gundersen, H.J., Jensen, E.B., 1987. The efficiency of systematic sampling in stereology and its prediction. J. Microsc. 147(Pt 3), 229–263.

Gundersen, H.J., Jensen, E.B., Kieu, K., Nielsen, J., 1999. The efficiency of systematic sampling in stereology--reconsidered. J. Microsc. 193(Pt 3), 199–211.

Harding, A., Halliday, G., Caine, D., Kril, J., 2000. Degeneration of anterior thalamic nuclei differentiates alcoholics with amnesia. Brain 123, 141–154.

Head, E., Powell, D., Gold, B.T., Schmitt, F.A., 2012. Alzheimer’s Disease in Down Syndrome. Eur J Neurodegener Dis 1(3), 353–364.

Hirai, T., Ohye, C., Nagaseki, Y., Matsumura, M., 1989. Cytometric analysis of the thalamic ventralis intermedius nucleus in humans. J. Neurophysiol. 61(3), 478–487.

Hof, P.R., Bouras, C., Perl, D.P., Sparks, D.L., Mehta, N., Morrison, J.H., 1995. Age-related distribution of neuropathologic changes in the cerebral cortex of patients with Down’s syndrome. Quantitative regional analysis and comparison with Alzheimer’s disease. Arch. Neurol. 52(4), 379–391.

Hornberger, M., Wong, S., Tan, R., Irish, M., Piguet, O., Kril, J., Hodges, J.R., Halliday, G., 2012. In vivo and post-mortem memory circuit integrity in frontotemporal dementia and Alzheimer’s disease. Brain 135(Pt 10), 3015–3025.

Hou, J., Riise, J., Pakkenberg, B., 2012. Application of immunohistochemistry in stereology for quantitative assessment of neural cell populations illustrated in the Gottingen minipig. PLoS One 7(8), e43556.

Jankowski, M.M., Ronnqvist, K.C., Tsanov, M., Vann, S.D., Wright, N.F., Erichsen, J.T., Aggleton, J.P., O’Mara, S.M., 2013. The anterior thalamus provides a subcortical circuit supporting memory and spatial navigation. Front Syst Neurosci 7, 45.

Jernigan, T.L., Bellugi, U., Sowell, E., Doherty, S., Hesselink, J.R., 1993. Cerebral morphologic distinctions between Williams and Down syndromes. Arch. Neurol. 50(2), 186–191.

Karlsen, A.S., Korbo, S., Uylings, H.B., Pakkenberg, B., 2014. A stereological study of the mediodorsal thalamic nucleus in Down syndrome. Neuroscience 279, 253–259.

Karlsen, A.S., Pakkenberg, B., 2011. Total numbers of neurons and glial cells in cortex and basal ganglia of aged brains with Down syndrome--a stereological study. Cereb. Cortex 21(11), 2519–2524.

Kemper, T.L., 1991. Down Syndrome, in: Peters, A., Jones, E.G. (Eds.), Cerebral Cortex: Normal and altered states of function. Plenum Press, pp. 511–526.

Krasuski, J.S., Alexander, G.E., Horwitz, B., Rapoport, S.I., Schapiro, M.B., 2002. Relation of medial temporal lobe volumes to age and memory function in nondemented adults with Down’s syndrome: implications for the prodromal phase of Alzheimer’s disease. Am. J. Psychiatry 159(1), 74–81.

Lakens, D., 2013. Calculating and reporting effect sizes to facilitate cumulative science: a practical primer for t-tests and ANOVAs. Front Psychol 4, 863.

Lyck, L., Jelsing, J., Jensen, P.S., Lambertsen, K.L., Pakkenberg, B., Finsen, B., 2006. Immunohistochemical visualization of neurons and specific glial cells for stereological application in the porcine neocortex. J. Neurosci. Methods 152(1-2), 229–242.

McGraw, K.O., Wong, S.P., 1992. A common language effect size statistic. Psychological Bulletin 111(2), 361.

Pelvig, D.P., Pakkenberg, H., Stark, A.K., Pakkenberg, B., 2008. Neocortical glial cell numbers in human brains. Neurobiol. Aging 29(11), 1754–1762.

Pinter, J.D., Brown, W.E., Eliez, S., Schmitt, J.E., Capone, G.T., Reiss, A.L., 2001. Amygdala and hippocampal volumes in children with Down syndrome: a high-resolution MRI study. Neurology 56(7), 972–974.

Raz, N., Torres, I.J., Briggs, S.D., Spencer, W.D., Thornton, A.E., Loken, W.J., Gunning, F.M., McQuain, J.D., Driesen, N.R., Acker, J.D., 1995. Selective neuroanatomic abnormalities in Down’s syndrome and their cognitive correlates: evidence from MRI morphometry. Neurology 45(2), 356–366.

Rub, U., Stratmann, K., Heinsen, H., Del Turco, D., Ghebremedhin, E., Seidel, K., den Dunnen, W., Korf, H.W., 2016. Hierarchical Distribution of the Tau Cytoskeletal Pathology in the Thalamus of Alzheimer’s Disease Patients. J Alzheimers Dis 49(4), 905–915.

Schimmel, M.S., Hammerman, C., Bromiker, R., Berger, I., 2006. Third ventricle enlargement among newborn infants with trisomy 21. Pediatrics 117(5), e928–931.

Tan, R.H., Wong, S., Kril, J.J., Piguet, O., Hornberger, M., Hodges, J.R., Halliday, G.M., 2014. Beyond the temporal pole: limbic memory circuit in the semantic variant of primary progressive aphasia. Brain 137(Pt 7), 2065–2076.

Teipel, S.J., Alexander, G.E., Schapiro, M.B., Moller, H.J., Rapoport, S.I., Hampel, H., 2004. Age-related cortical grey matter reductions in non-demented Down’s syndrome adults determined by MRI with voxel-based morphometry. Brain 127(Pt 4), 811–824.

Tsivilis, D., Vann, S.D., Denby, C., Roberts, N., Mayes, A.R., Montaldi, D., Aggleton, J.P., 2008. A disproportionate role for the fornix and mammillary bodies in recall versus recognition memory. Nat. Neurosci. 11(7), 834–842.

Van der Werf, Y.D., Jolles, J., Witter, M.P., Uylings, H.B., 2003. Contributions of thalamic nuclei to declarative memory functioning. Cortex 39(4-5), 1047–1062.

Vann, S.D., Tsivilis, D., Denby, C.E., Quamme, J.R., Yonelinas, A.P., Aggleton, J.P., Montaldi, D., Mayes, A.R., 2009. Impaired recollection but spared familiarity in patients with extended hippocampal system damage revealed by 3 convergent methods. Proceedings of the Academy of Natural Sciences 106(13), 5442–5447.

Vicari, S., Bellucci, S., Carlesimo, G.A., 2000. Implicit and explicit memory: a functional dissociation in persons with Down syndrome. Neuropsychologia 38(3), 240–251.

Weis, S., Weber, G., Neuhold, A., Rett, A., 1991. Down syndrome: MR quantification of brain structures and comparison with normal control subjects. AJNR. American journal of neuroradiology 12(6), 1207–1211.

West, M.J., Slomianka, L., Gundersen, H.J., 1991. Unbiased stereological estimation of the total number of neurons in thesubdivisions of the rat hippocampus using the optical fractionator. Anat Rec 231(4), 482–497.

Wu, J., Morris, J.K., 2013. The population prevalence of Down’s syndrome in England and Wales in 2011. Eur J Hum Genet 21(9), 1016–1019.

Xuereb, J.H., Perry, R.H., Candy, J.M., Perry, E.K., Marshall, E., Bonham, J.R., 1991. Nerve cell loss in the thalamus in Alzheimer’s disease and Parkinson’s disease. Brain 114 (Pt 3), 1363–1379.

Zigman, W.B., Lott, I.T., 2007. Alzheimer’s disease in Down syndrome: neurobiology and risk. Ment Retard Dev Disabil Res Rev 13(3), 237–246.

